# TES-1/Tes protects junctional actin networks under tension from self-injury during epidermal morphogenesis in the *C. elegans* embryo

**DOI:** 10.1101/2021.09.30.462556

**Authors:** Allison M. Lynch, Bethany G. Lucas, Jonathan D. Winkelman, Sterling C.T. Martin, Samuel D. Block, Anjon Audhya, Margaret L. Gardel, Jeff Hardin

## Abstract

During embryonic morphogenesis, the integrity of epithelial tissues depends on the ability of cells in tissue sheets to undergo rapid changes in cell shape while preventing self-injury to junctional actin networks. LIM domain-containing repeat (LCR) proteins are recruited to sites of strained actin filaments in cultured cells [1–3], and are therefore promising candidates for mediating self-healing of actin networks, but whether they play similar roles in living organisms has not been determined. Here, we establish roles for *Caenorhabditis elegans* TES-1/Tes, an actin-binding LCR protein present at apical junctions, during epithelial morphogenesis. TES-1::GFP is recruited to apical junctions during embryonic elongation, when junctions are under tension; in embryos in which stochastic failure of cell elongation occurs, TES-1 is only strongly recruited to junctions in cells that successfully elongate, and recruitment is severely compromised in genetic backgrounds in which cell shape changes do not successfully occur. *tes-1* mutant embryos display junctional F-actin defects, and loss of TES-1 strongly enhances tension-dependent injury of junctional actin networks in hypomorphic mutant backgrounds for CCC components, suggesting that TES-1 helps to prevent self-injury of junctional actin networks during rapid cell shape change. Consistent with such role, a fragment of TES-1 containing its LIM domains localizes to stress fiber strain sites (SFSS) in cultured vertebrate cells. Together, these data establish TES-1 as a tension-sensitive stabilizer of the junctional actin cytoskeleton during embryonic morphogenesis.

## Introduction

Embryonic tissues require epithelial cell-cell adhesions that are both dynamic and strong. On the one hand, they must be dynamic, as cells rearrange and change shape to allow for the complex processes of morphogenesis, but on the other, cell-cell adhesions must be able to withstand contractile forces that threaten tissue integrity [4–7]. Thus, identifying factors that modulate junctional integrity is important for understanding embryonic morphogenesis. Here, we describe a novel modulatory role for the *C. elegans* Testin/Tes ortholog, TES-1, at cell-cell junctions.

Vertebrate Tes/testin (hereafter Tes) has an N-terminal CR domain, a PET (Prickle, Espinas and Testin) domain, and three C-terminal LIM (Lin11, Isl-1 & Mec-3) domains. Biochemical and structure-function analyses have suggested that the N terminus of Tes allows association with the actin cytoskeleton [8, 9]. The LIM domains appear to allow association with heterologous binding partners [10, 11], may be involved in mediating intracellular inhibition of Tes [12] and the PET domain may allow homodimerization of Tes via interaction with the LIM1-2 region [13].

Tes has been implicated in several actin-dependent processes. In cultured cells, Tes localizes to focal adhesions, integrin-based attachment sites linking intracellular actin stress fibers to sites of attachment to the extracellular matrix (ECM) [1, 8, 9, 14]. Tes also appears to associate with cell-cell adhesions. Tes localizes to spot-like cell-cell contacts [10, 15], where it colocalizes with cadherin/catenin complex (CCC) components [10]. Vertebrate zyxin can interact with Tes in vitro [9, 11] and also localizes to adherens junctions [8, 9, 15–22]. Zyxin and other LIM domain proteins preferentially localize to natural rupture sites in bundled F-actin networks in cultured cells subjected to tension, where they are thought to allow rapid healing of ruptured bundles [23–25]; reviewed in [1]. In cultured cells the LIM domains of zyxin, Tes, and other LIM domain proteins are sufficient for this response [2].

While previous work has examined the role of Tes in establishing and maintaining actin networks under stress in cultured cells, particularly at sites of cell-ECM attachment, roles for Tes have not been established at sites of epithelial cell-cell adhesion in an intact, developing organism. *C. elegans* epidermal morphogenesis provides a convenient system for studying the roles of proteins that modulate epithelial cell-cell adhesion. Epidermal morphogenetic movements that require the CCC include (1) ventral enclosure, during which the embryo is encased in an epidermal monolayer [26] and new epidermal cell-cell junctions are formed [27]; and (2) the early phase of elongation, which involves a coordinated actomyosin-mediated contraction of the embryo, primarily driven by lateral (seam) epidermal cells [28, 29]; reviewed in [30, 31]. These contractile forces exert substantial tension on cell-cell junctions; the forces of contraction are transmitted via circumferential filament bundles (CFBs), large bundles of F-actin anchored at epidermal cell-cell junctions [28, 32]. Anchorage of CFBs depends on the core components of the CCC: HMR-1/cadherin, HMP-2/β-catenin, and HMP-1/α-catenin [27]. Reduced function of core CCC components coupled with removal of other adherens junctional proteins leads to catastrophic morphogenetic failure, predominantly during embryonic elongation [33–36]. Here, we describe the roles of TES-1/Tes, in preventing “self-injury” of the junctional proximal actin network that maintains the connection between the CCC and CFBs during periods of mechanical stress in the developing *C. elegans* epidermis.

## Results and Discussion

We previously conducted a genome-wide RNAi screen in a sensitized HMP-1/α-catenin background, *hmp-1(fe4)*, and uncovered modulators of cell adhesion in *C. elegans* during morphogenesis [35]. In our initial screen, we identified a gene on chromosome IV, which when knocked down, potently enhanced the penetrance and severity of the *hmp-1(fe4)* phenotype [35] (Supplemental Video 1). Previously named TAG-224 (Temporarily Assigned Gene 224), we renamed the protein TES-1 because of its significant homology to the vertebrate protein Tes after examination of the predicted protein sequence using BLAST and ClustalW. ClustalW analysis indicates that the two proteins are approximately 35% identical and 64% similar. Pfam analysis shows both proteins have a similar domain structure: an N-terminal PET domain followed by three C-terminal LIM domains (Fig. 1A).

**Figure 1.**
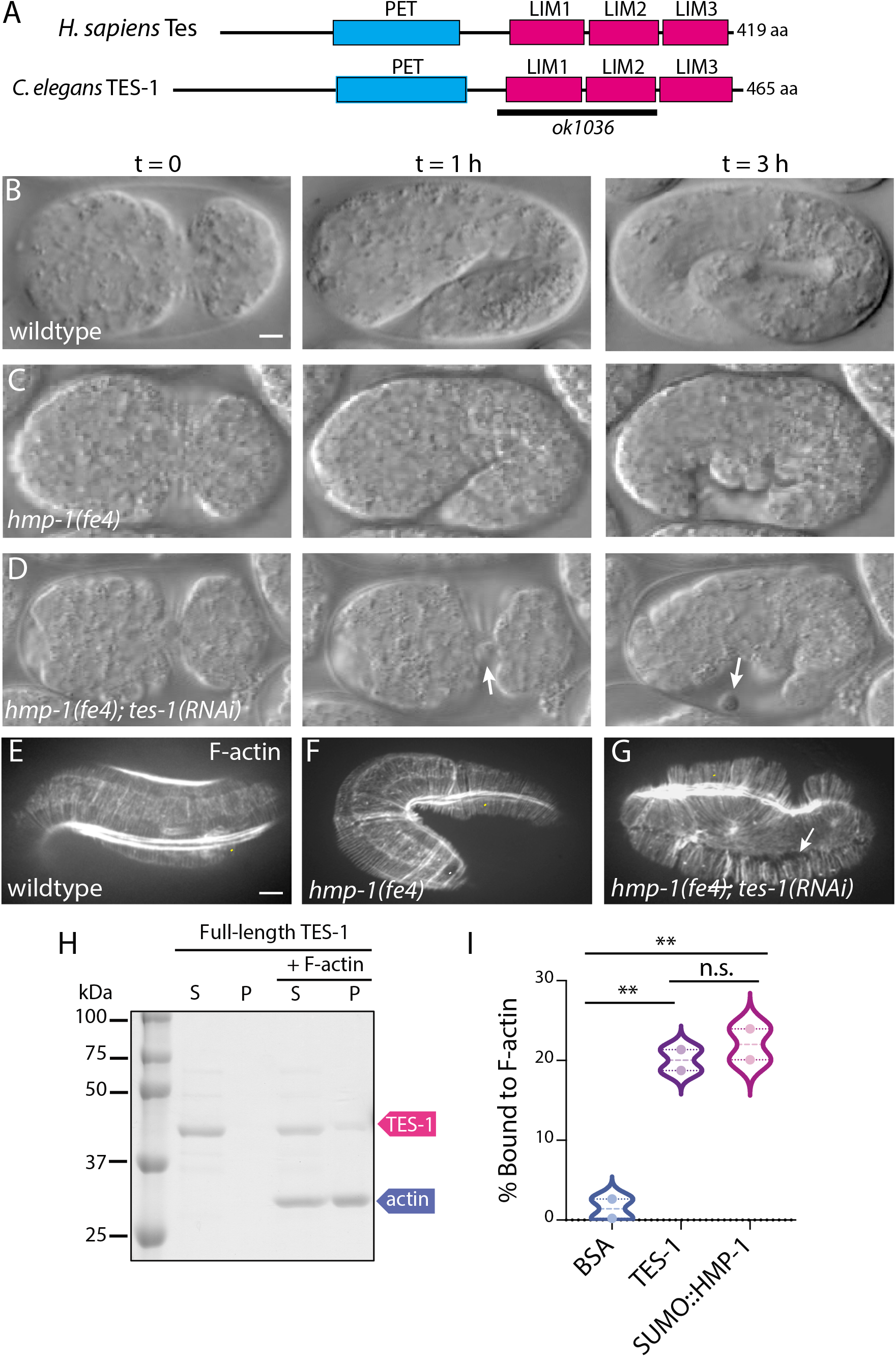
TES-1 loss enhances phenotypes in hypomorphic CCC backgrounds. (A) Protein domain maps of *C. elegans* TES-1 and human Tes. TES-1 and Tes both contain N-terminal Prickle, Espinas, Testin (PET) domains and three C-terminal Lin-11, Isl-1, Mec-3 (LIM) domains. The *tes-1(ok1036)* allele removes LIM1-2 along with some intronic sequence and introduces a frameshift into the remainder of the coding region. (B-D) *tes-1(RNAi)* enhances the severity of morphogenetic defects in *hmp-1(fe4)* embryos. (B) Wild-type embryo imaged using Nomarski microscopy. (C) *hmp-1(fe4)* embryo; bulges become apparent during embryonic elongation (t = 3 hr). (D) In *hmp-1(fe4);tes-1(RNAi)* embryos, cells leak out of the ventral midline (t = 1 hr), and all embryos die with sever elongation defects (t = 3 hr). (E-G) *tes-1(RNAi)* enhances the severity of actin defects in *hmp-1(fe4)* embryos. (E) Wild-type embryos maintain a population of junctional proximal actin along cell borders (white arrow) and dorsal and ventral epidermal cells in elongated embryos contain circumferential actin filament bundles (CFBs) that are evenly spaced. Bright signal is muscle, denoted by yellow arrowheads. (F) *hmp-1(fe4)* embryos also typically maintain junctional proximal actin (white arrow); however, their CFBs are less evenly spaced, and sometimes clump together (white arrowhead). (G) *hmp-1(fe4); tes-1(RNAi)* embryos display clumping of CFBs, and also a complete lack of junctional proximal actin. CFBs appear to have been torn away from the junction, leaving bare zones devoid of Factin (white arrow). Scale bar = 5 μm. (H) TES-1 binds to F-actin in an actin co-sedimentation assay. Full-length TES-1 remains in the supernatant fraction (S) when incubated without F-actin. However, TES-1 is detected in the pellet fraction (P) when incubated with 5 μM F-actin. (I) Quantification of TES-1 found in the pellet after incubation with F-actin. Bovine Serum Albumin (BSA) served as a negative control and SUMO:HMP-1 as a positive control. TES-1 bound to Factin significantly more than did BSA (two replicates; **p < = 0.01, unpaired Student’s T test).

### *TES-1 functionally interacts with hmp-1/α-catenin at the* C. elegans *apical junction*

All *hmp-1(fe4); tes-1*(RNAi) embryos arrest during the elongation stage of morphogenesis with junctional actin defects that suggest a requirement for TES-1 during developmental stages requiring strong cell-cell adhesions (Fig. 1B-D). In 26 % of *hmp-1(fe4); tes-1*(RNAi) embryos (6 of 23 embryos examined via 4d microscopy) individual cells leaked out of the ventral midline, compared with 0% of *hmp-1(fe4)* homozygotes (0 of 22 embryos examined; significantly different, Fisher’s exact test, p = 0.02), suggesting that nascent junctions also have a more stringent requirement for TES-1 in this sensitized background (Fig. 1D, arrow). Phalloidin staining demonstrated that *tes-1(RNAi)* exacerbated junctional proximal actin defects in a *hmp-1(fe4)* background (Fig. 1E-G). We confirmed the efficacy of *tes-1* RNAi and the synergistic lethality resulting from TES-1 depletion in *hmp-1(fe4)* embryos by crossing a deletion allele, *tes-1(ok1036)*, into *hmp-1(fe4)* worms: double homozygotes exhibit 93.7% lethality and elongation arrest (n = 516 embryos examined).

We next confirmed that, like vertebrate Tes [8, 9], TES-1 directly binds F-actin by performing actin cosedimentation assays using recombinant TES-1 protein and found that TES-1 cosediments with F-actin (Fig. 1H). The extent of cosedimentation of TES-1 with F-actin is statistically indistinguishable from another well characterized junctional actin-binding protein in *C. elegans*, HMP-1/α-catenin [37] (Fig. 1I).

### TES-1 localizes to apical junctions in the embryonic epidermis

To assess the expression pattern and subcellular localization of TES-1, we constructed a translational fusion protein containing the entire genomic sequence of *tes-1* fused to GFP driven by its endogenous promoter (Fig. 2). In early embryos, TES-1::GFP is visible in epidermal cells, where its location is exclusively cytoplasmic. At the 2-fold stage of elongation, TES-1::GFP puncta begin to accumulate at sites of cell-cell contact (Fig. 2B). These clusters expand and become more evenly distributed along cell borders as elongation continues (Fig. 2C); junctional TES-1 signal likewise increases significantly after the two-fold stage of elongation (Fig. 2H). Strikingly, TES-1::GFP is maintained at seam-dorsal and seam-ventral, but not seam-seam borders (Fig. 2D). Importantly, TES-1::GFP rescues lethality seen in *ok1036/+; fe4* embryos. *ok1036/+;fe4/fe4* worms are extremely difficult to maintain due to *fe4* maternal effect; progeny of such worms exhibit 80% lethality (n = 20 embryos scored) and the addition of extrachromosomal TES-1::GFP reduces this lethality to 38% (n = 92 embryos scored). Significantly, *ok1036/ok1036;fe4/fe4* worms can develop to adulthood, but only if they express *tes-1::gfp*, indicating the GFP is functional.

**Figure 2.**
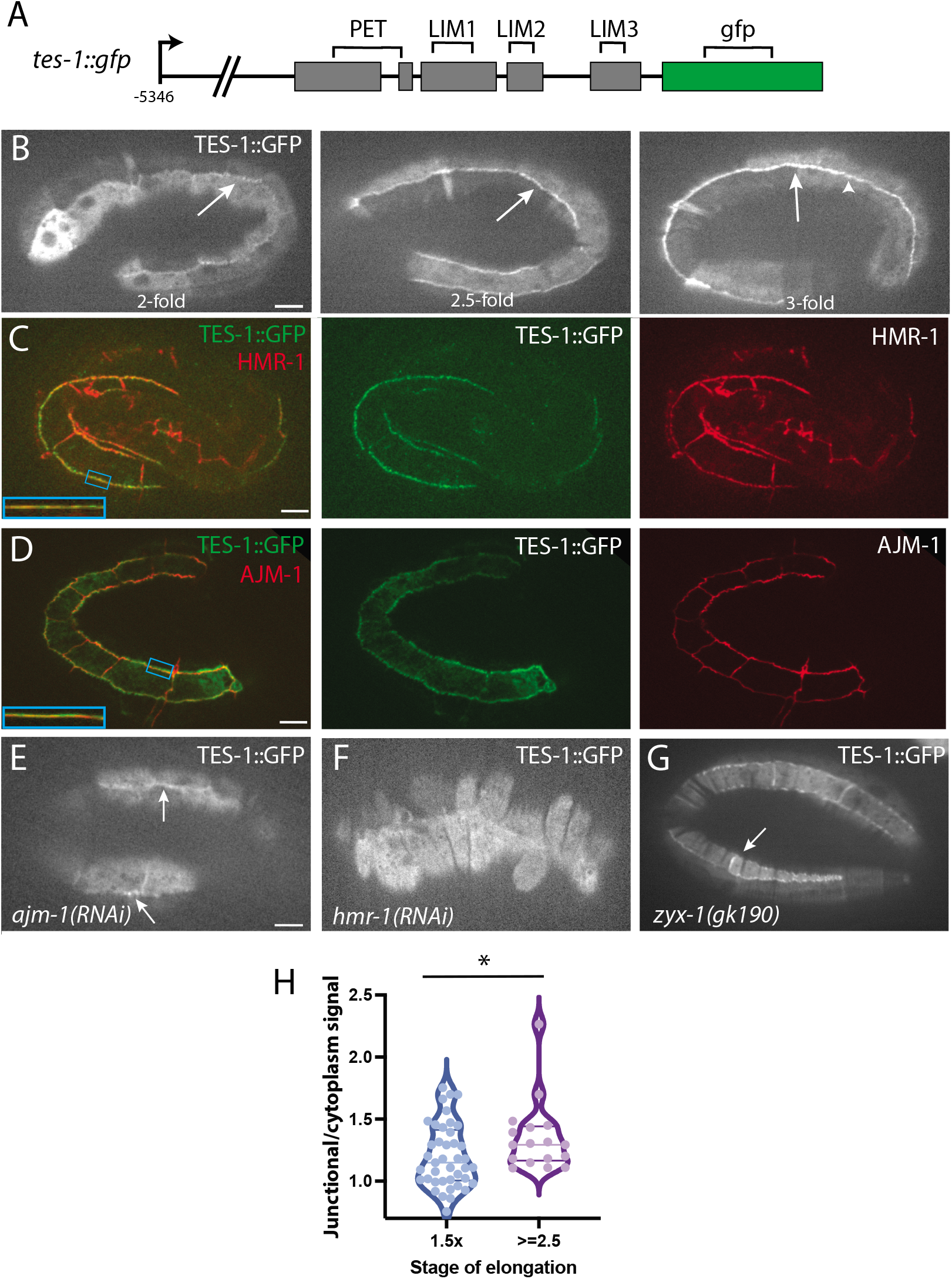
TES-1 localizes to sites of cell-cell attachment during embryonic elongation. (A) A schematic of the full-length TES-1::GFP used in this study, along with TES-1::GFP is driven by its full-length endogenous promoter and contains a C-terminal GFP. (B) Expression of TES-1::GFP is initially cytoplasmic, but as elongation proceeds, TES-1::GFP localizes to cellcell junctions, first in puncta and then strongly to seam-dorsal and seam-ventral boundaries. Scale bar = 5 μm. (C-D) TES-1::GFP co-localizes with HMR-1 but not AJM-1. Insets show magnifications of boxed regions. While HMR-1 and TES-1::GFP largely co-localize, TES-1::GFP and AJM-1 do not. (E) *ajm-(RNAi)* does not influence the ability of TES-1::GFP to localize to junctions (arrows). Scale bar = 5 μm. (F) *hmr-1(RNAi)* completely prevents TES-1::GFP localization at junctions (arrow); however, the embryos never elongate to the point at which TES-1::GFP normally localizes. (G) *zyx-1(gk190); tes-1::gfp* embryos show persistent seam-seam junctional localization in addition to the normal seam-dorsal and seam-ventral localization (arrow). (H) TES-1 is recruited to junctions during elongation. Junctional/cytoplasmic ratio of TES-1::GFP at 1.5-fold and >=2.5-fold. *, p < 0.05 (Mann-Whitney U test).

To determine whether TES-1::GFP colocalizes with adhesion complexes, we performed immunostaining experiments on embryos expressing TES-1::GFP. The apicobasal distribution of TES-1 indicates that it colocalizes predominatly with the cadherin/catenin complex vs. the DLG-1/AJM-1 complex. Embryos co-immunostained for HMR-1/cadherin and GFP display substantial overlap of HMR-1 and TES-1::GFP (Fig. 2C), wheras there is much less overlap with AJM-1, a component of the DLG-1 (Discs Large)/AJM-1 complex, which is basal to the CCC (Fig. 2D). Partial localization of Tes with the CCC has likewise previously been reported in cultured vertebrate cells [10]. Although one study has reported that vertebrate α-catenin and Tes can be coimmunoprecipitated [38], we have been unable to coimmunoprecipitate TES with *C. elegans* CCC components in a generalized proteomics approach [39] or in directed coIP experiments (Fig. S2), suggesting that the interaction of TES-1 with the *C. elegans* CCC is indirect.

To address the role of junctional components in localizing Tes in living embryos, we performed knockdown experiments in *tes-1::gfp* transgenic embryos followed by confocal microscopy. *ajm-1*(RNAi) in embryos expressing TES-1::GFP does not prevent localization of TES-1::GFP to junctions (Fig. 2E). In these embryos, however, TES-1::GFP foci do not spread to form a continuous, intense band, which may reflect the failure of *ajm-1(RNAi)* embryos to elongate successfully (see below). In contrast, the effects of loss of function of *hmr-1* /cadherin on TES-1::GFP localization were more severe. In *hmr-1*(RNAi) embryos TES-1::GFP failed to be recruited to junctions (Fig. 2F).

Based on previous studies of vertebrate homologues [9, 11], we next assessed the effects of loss of ZYX-1/zyxin on TES-1::GFP junctional localization, since ZYX-1 translational fusions have previously been reported to be weakly expressed in epidermal cells in embryos [40]. While vertebrate Tes can physically interact with zyxin [9, 11] and we were able to coIP TES-1 and ZTX-1 (Fig. S3A), we were only able to detect a very weak, substoichiometric interaction between TES-1 and ZYX-1 via coIP and pulldown of bacterial expressed proteins (Fig. S3B). When we crossed the strain expressing *tes-1::gfp* into *zyx-1(gk190)* worms, we occasionally saw TES-1::GFP localized to seam-seam junctions in addition to seam-dorsal and seam-ventral junctions in resulting progeny at the 3-4-fold stage (Fig. 2G; 4/13 embryos examined; 0/8 embryos examined displayed seam:seam localization in the transgenic strain alone; not significant, Fisher’s Exact test, p = 0.27), suggesting that ZYX-1 contributes weakly at best to the restriction of TES-1 to seam-dorsal and seam-ventral boundaries. This result is consistent with experiments in vertebrates, which show that while depletion of Zyxin can reduce the amount of Tes at focal adhesions [9], Tes can still localize independently of Zyxin [11].

### TES-1 requires its PET and LIM2-3 domains

To our knowledge, the roles of specific regions of a Tes family member have not been assessed in a living organism. To identify which subdomains are required for junctional targeting and function of TES-1 we analyzed the expression pattern of GFP deletion constructs and analyzed the ability of each construct to rescue embryonic viability in offspring from *hmp-1(fe4)/+; tes-1(ok1036)/+* mothers (Fig. 3). Unlike full-length TES-1::GFP, TES-1ΔPET::GFP localized along all seam cell borders in the epidermis, including seam-seam borders, as well as CFBs (Fig. 3B). Deletion of LIM1 (Fig. 3C) or LIM2 (Fig. 3D) both perturbed junctional localization similarly: each localized sporadically to epidermal junctions, including some seam-seam junctions. However, there was also localization at what appeared to be actin-containing structures in epidermal cells. Deletion of LIM3 rendered the GFP entirely cytoplasmic (Fig. 3E). Deletion of all three LIM domains simultaneously resulted in GFP localization along structures that appear to be CFBs (Fig. S3F). This result is consistent with work on vertebrate Tes, which can coimmunoprecipitate actin [9] and localize via its N terminus [13, 38]. Because full-length TES-1::GFP localizes to cell-cell junctions, this result suggests that the latent ability of TES-1 to bind to CFBs is not normally manifest when other subdomains of TES-1 can bind their normal targets.

**Figure 3.**
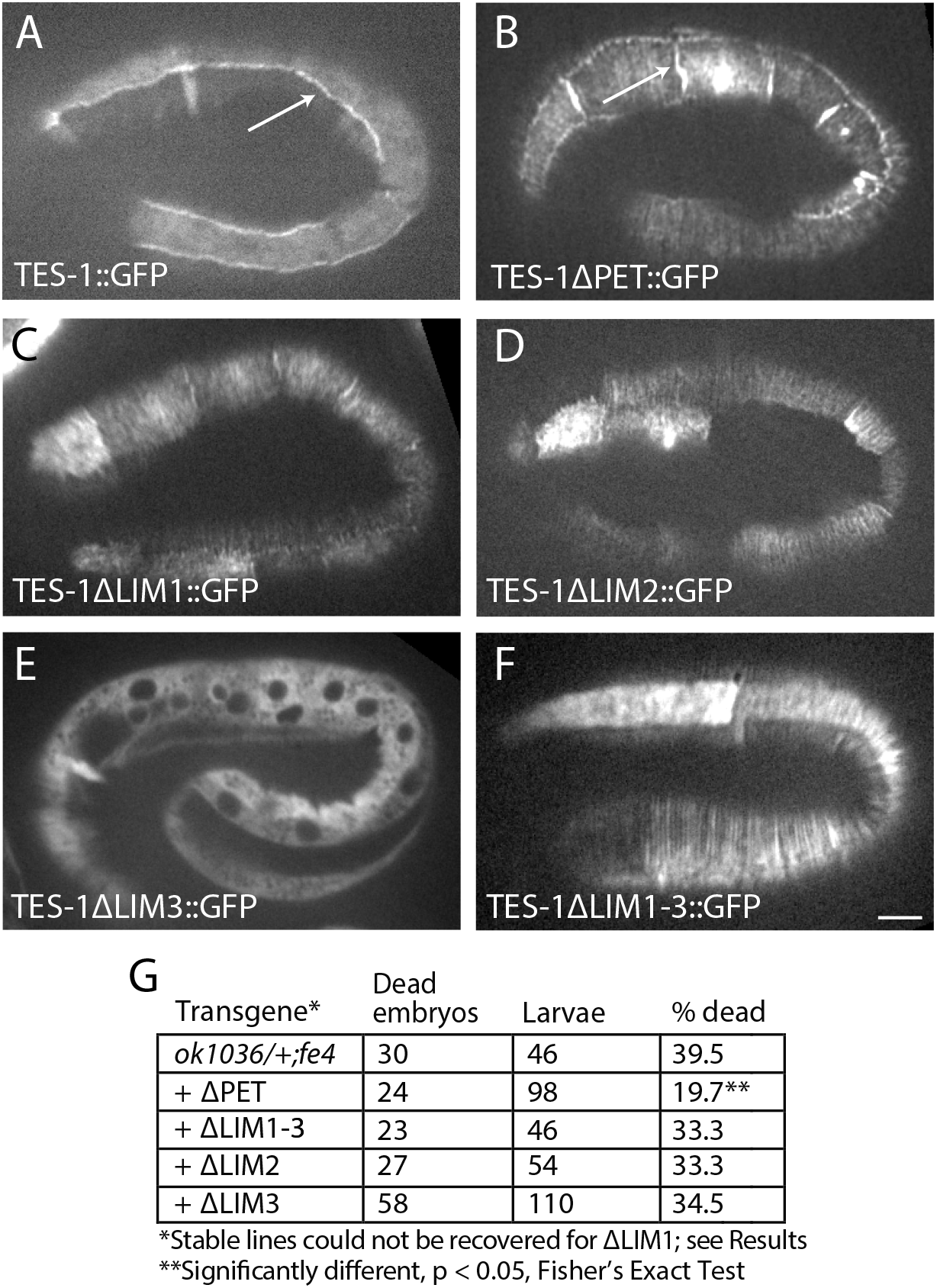
TES-1 localization requires its PET and LIM2-3 domains. For relevant domains of TES-1, see Figure 1A. (A) Full-length TES-1::GFP localizes to dorsal:seam and ventral:seam cell boundaries in the epidermis (arrow). (B) Unlike full-length TES-1::GFP, TES-1 ΔPET::GFP localizes along all seam cell borders in the epidermis, including seam-seam borders (arrows). Deletion of LIM1 (C) or LIM2 (D) both perturb junctional localization similarly: each localizes sporadically to epidermal junctions, including some seam-seam junctions. However, there is also localization at what appeared to be actin-containing structures in epidermal cells. (E) Deletion of LIM3 rendered the GFP entirely cytoplasmic. (F) Deletion of all three LIM domains simultaneously resulted in GFP localization along structures that appear to be CFBs. (G) Rescue of embryonic lethality in progeny of *tes-1(ok1036)/+;hmp-1(fe4)/hmp-1(fe4)* hermaphrodites. * = significantly different from non-transgenic animals (p < 0.05, Fisher’s exact test).

Due to maternal effects and gonadal defects, assessing synergistic lethality of *tes-1::gfp* deletion constructs in *hmp-1(fe4)* homozygous mothers proved challenging. We therefore tested for the ability of subdomains of TES-1 to rescue synergistic lethality in *ok1036/ok1036; hmp-1(fe4)/+* embryos (Fig. 3G). TES-1ΔPET could rescue some embryonic lethality in this genetic background, but progeny had numerous defects, including germline malformations, protruding vulvae, and sterility. TES-1ΔLIM1-3, TES-1ΔLIM2, and TES-1ΔLIM2 were unable to rescue the 39% lethality observed among progeny of *ok1036/ok1036; hmp-1(fe4)/+* mothers. among progeny of *tes-1(ok1036); hmp-1(fe4)* embryos. Interestingly, while lines harboring *tes-1ΔLIM1::GFP* could not be obtained, occasional *tes-1(ok1036); hmp-1(fe4); tes-1 ΔLIM1::GFP* embryos were able to grow to adulthood, but these adults were sterile. Overall, these results indicate that the LIM domains of TES-1 are crucial for *tes-1* function during morphogenesis, but that LIM1 may be less important, and are consistent with the observation that the distribution of TES-1ΔPET::GFP and TES-1ΔLIM1::GFP appears most similar to full-length TES-1::GFP.

While overall the deletion analysis indicates that the LIM domains are crucial for junctional targeting of TES-1, the difference in localization pattern of the ΔLIM3 and ΔLIM1-3 is curious. Recently, it has been suggested that the LIM1-2 domain of vertebrate TES can engage in both heterophilic binding to proteins such as zyxin and homophilic dimerization via interaction with the PET domain of Tes [38]. Homodimerization of αE-catenin drives it away from adherens junctions [41, 42]. While it is not currently known if homodimeric Tes is sequestered away from cell adhesion sites in a similar way, if it is this might explain the cytoplasmic accumulation of TES-1 ΔLIM3::GFP in *C. elegans*. Deletion of LIM3 might favor homodimerization over heterophilic interactions of TES-1 with other binding partners.

### TES-1 localizes to junctions in a tension-dependent manner

Tes is required for the maintenance of stress fibers in cultured vertebrate cells [43], and accumulates at “focal adherens junctions”, spot-like foci of cell-cell adhesion, in human vascular endothelial cells [10]. These data suggest that Tes might play tension-dependent roles in organizing the actin network at adherens junctions in epithelia during embryonic morphogenesis. During elongation of the *C. elegans* embryo, a coordinated change in the shape of epidermal cells drives elongation of the embryo to approximately 4-fold its original length [28]. The CCC anchors CFBs at junctions – specifically seam-ventral and seam-dorsal junctions – during this time, when the contractile forces driving elongation result in elevated tension at these junctional boundaries [27, 32, 44–46]. Given the localization of TES-1, it is a good candidate to stabilize junctional actin networks during embryonic elongation.

Because *hmr-1* /cadherin, *hmp-1/α-catenin*, and *hmp-2/β-catenin* homozygous null mutant embryos fail to progress past the two-fold stage of elongation, we could not assess whether disruption of TES-1:: GFP recruitment to junctions is due primarily to physical absence of CCC components or because of the pre-elongation death of the embryos. In order to adjudicate between these possibilities we examined *hmp-1(fe4)* embryos expressing TES-1::GFP. The *fe4* lesion causes weaker binding of F-actin by HMP-1 and leads to less stable junctions [47]. *hmp-1(fe4)* embryos display a variable phenotype (Fig. 4A); while some embryos fail to elongate appreciably, other embryos extend to the 2-fold stage of elongation (Fig. 4B). We found that TES-1::GFP did not localize to junctions in *hmp-1(fe4)* embryos that failed to elongate past 1.5-fold (5 of 5 embryos imaged via spinning disc confocal microscopy), even in embryos that survived and hatched. However, TES-1::GFP did localize to junctions in *hmp-1(fe4)* embryos that elongated to at least 2-fold their original length (3 of 3 embryos examined; significantly different; Fisher’s exact test, p = 0.018). The correlation between the extent of elongation of *fe4* embryos and the normal TES-1::GFP localization patterns identified previously suggests that TES-1 is only recruited to junctions that resemble those in normal embryos at the 2-fold stage.

**Figure 4.**
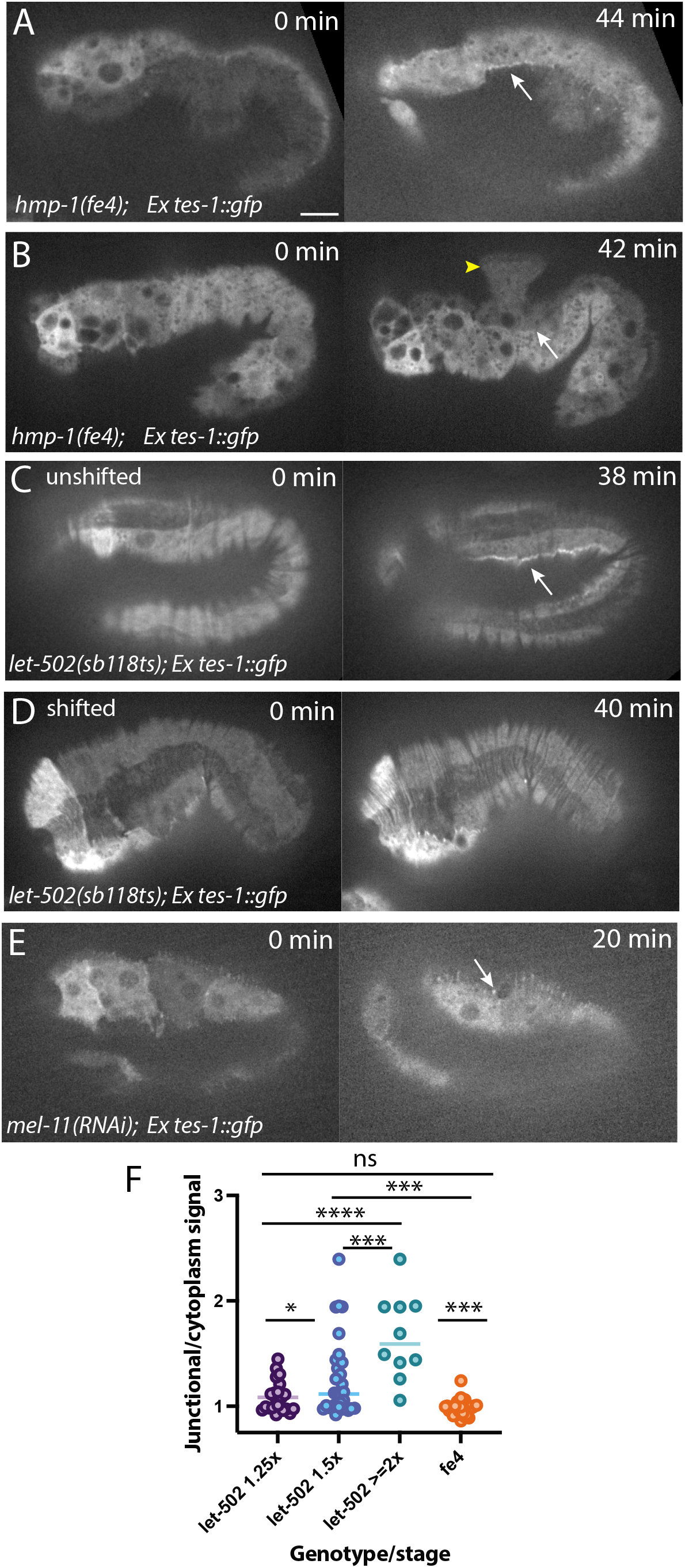
TES-1::GFP localizes to junctions in a tension-dependent manner. (A) In *hmp-1(fe4)* embryos that successfully elongate to two-fold, TES-1::GFP accumulates along seam cell junctions (white arrow). (B) In *hmp-1(fe4)* embryos that do not elongate past 1.5-fold before failing, TES-1::GFP does not localize to junctions, instead remaining entirely cytoplasmic. (C) In *let-502(sb118ts); tes-1::gfp* embryos reared at the permissive temperature (“unshifted”), development is normal and TES-1::GFP localizes to junctions as in wildtype. (D) In temperature-shifted embryos, the LET-502 protein is inactivated, embryos fail to elongate, and TES-1::GFP never accumulates along epidermal junctions. (E) In *mel-11(RNAi); tes-1::gfp* embryos, the embryos elongate normally, and TES-1::GFP junctional localization is not impacted. (F) Upon depletion of *mel-11*, the worms elongate past their normal four-fold length. TES-1::GFP localizes to junctions, however it appears as though it is being pulled away from junctions in long extensions from epidermal cell borders. Scale bars = 5 μm.

To examine whether junctional tension affects the ability of TES-1::GFP to localize, we introduced the full-length TES-1::GFP into *let-502(sb118)* worms (Fig. 4C-D). Loss of LET-502/Rho kinase reduces actomyosin contractility in the epidermis and prevents elongation of *C. elegans* embryos. *let-502(sb118)* is a temperature-sensitive allele; when *let-502(sb1180); tes-1::gfp* embryos were imaged at permissive temperatures, TES-1::GFP localized to junctions in a wild-type manner (Fig. 4C). However, when these embryos were reared at the restrictive temperature (25°C), TES-1::GFP remained entirely cytoplasmic in embryos that failed to elongate (Fig. 3D). We also attempted the converse experiment: loss of MEL-11/myosin phosphatase function results in excessively elongated embryos due to greater than normal epidermal contractility [45, 46]. However, adhesion complexes undergo changes in morphology that make this converse experiment difficult to interpret. In MEL-11-depleted embryos, the initially continuous distribution of junctional TES-1::GFP was progressively lost, as TES-1::GFP became fragmented and pulled away from junctions into linear arrays (Fig. 4E,F). One possibility consistent with this result is that the excessive tension that develops in a *mel-11* loss-of-function background leads to collapse of junctional proximal actin around CFB insertion sites, including associated TES-1.

### TES-1 regulates actin networks during elongation

We next assessed why loss of TES-1 might enhance the *hmp-1(fe4)* phenotype. LIM domains proteins, including Tes and zyxin, are recruited to “stress fiber strain sites”, bundled Factin networks in cultured cells subjected to strain, where they are thought to allow rapid healing of damaged bundles [2, 23–25]. We reasoned that since under normal circumstances TES-1 colocalizes with the CCC, TES-1 could similarly help to stabilize junctional-proximal actin subject to strain during periods of mechanical stress at adherens junctions during elongation. Consistent with this possibility, when we examined F-actin organization in *tes-1(ok1036)* homozygous embryos via phalloidin staining, we found defects not present in wild-type embryos (Fig. 5C-F). As seen in Fig. 5D, the majority of *tes-1(ok1036)* embryos display decreased junctional-proximal actin. Additionally, we also observed more severe phenotypes, including gaps between CFBs, CFB collapse, and complete loss of preserved junctional-proximal actin (Fig. 5E). These defects are consistent with TES-1 preventing damage to junctional actin networks subjected to high strain during embryonic elongation.

**Figure 5.**
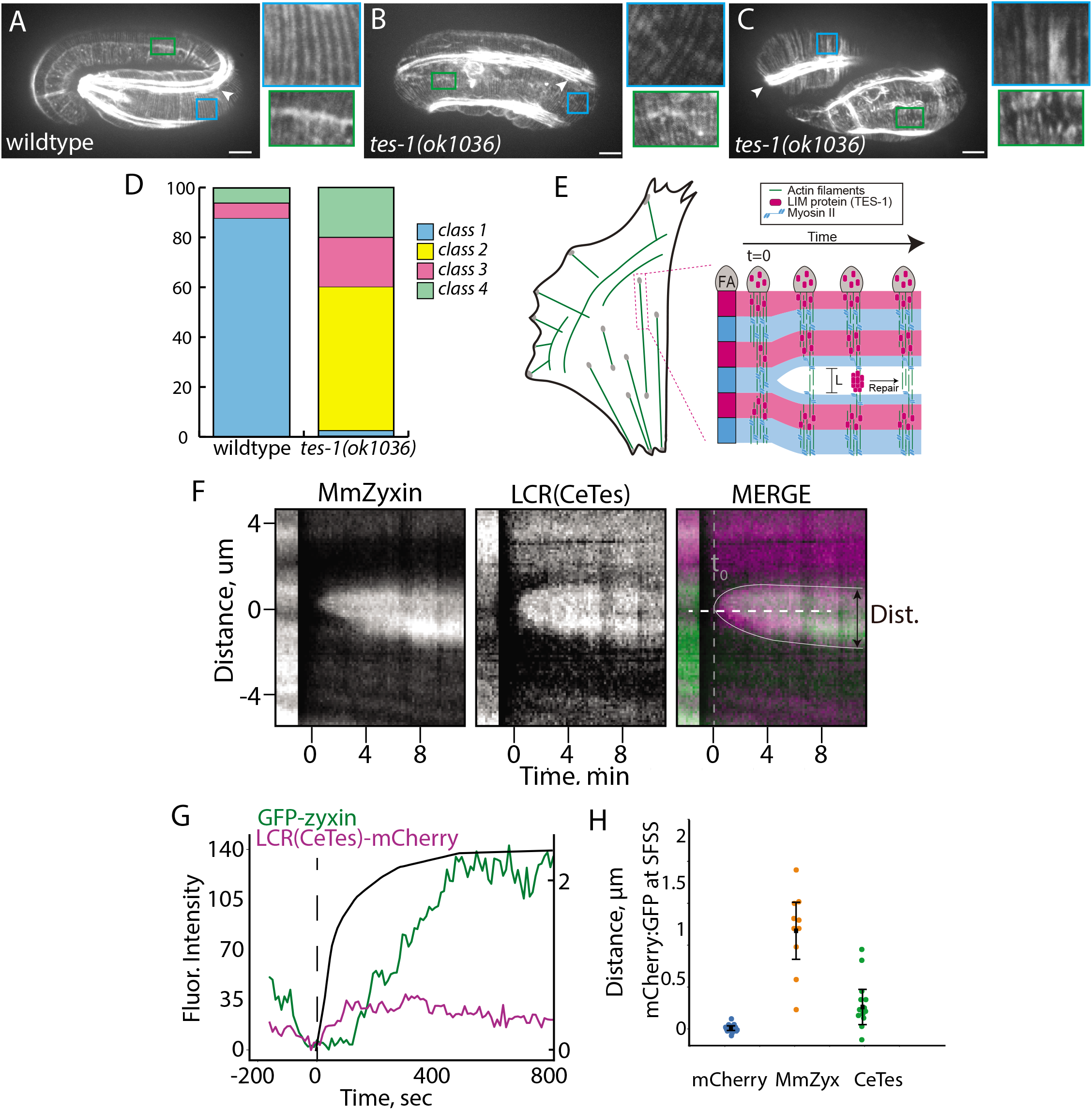
TES-1 regulates actin networks in vivo and is recruited to strained actin filaments. (A-D) Fixed and phalloidin stained embryos. Bright staining is muscle (arrowhead). Scale bar is 5 μm.(A) Wild-type embryos exhibit parallel circumferential filament bundles (CFBs, blue box inset) and retain junctional-proximal actin (green box inset). (B) Approximately half the *tes-1(ok1036)* embryos exhibit reduced junctional-proximal actin although CFB organization looks normal. (C) *ok1036* embryos also exhibit more severe phenotypes including gaps and clumping of CFBs (blue box) and a complete loss of junctional-proximal actin (green box). (D) Quantification of phalloidin staining phenotypes. Class 1 embryos have normal CFBs and junctional-proximal actin. Class 2 embryos have reduced junctional-proximal actin. Class 3 embryos have reduced junctional-proximal actin and CFB organization defects and Class 4 embryos have no retained junctional-proximal actin and CFB organization defects. (E-H) Recruitment of the TES-1 LCR::mCherry to stress fiber strain sites (SFSS) in transfected mouse embryonic fibroblasts. (E) Schematic of the experiment (adapted from [2]). Irradiated stress fibers under strain recruit LCR proteins, which are visible as a zone of recruitment in kymographs. (F) Representative kymographs of laser-induced recruitment of the TES-1 LCR::mCherry and mouse GFP::Zyxin to SFSS. White dashed and gray solid lines indicate where fluorescence and distance were measured. Dashed gray vertical line indicates to, when strain is first observed (H) Quantification of GFP and mCherry accumulation over time in the kymograph from (G). (H) LCR mCherry/GFP ratio for MmZyx (red) and TES-1 LCR (green). TES-1 LCR accumulates markedly (p=0.23, n>10; Student’s T-test, equal variance not assumed) but to a lesser extent than MmZyx; error bars indicate 95% confidence intervals.

Mammalian LIM domain proteins are recruited to strained actin networks via their LIM domain-containing region [2, 3, 24]. Since removal of the LIM domains of TES-1 results in loss of recruitment to junctional actin networks in the epidermis, we tested whether the LCR of TES-1 behaves similarly. When transfected into mouse embryonic fibroblasts, TES-1(LIM1-3)::mCherry is recruited to damaged actin bundles, indicating that it behaves in a manner similar to other LIM domain proteins in this assay. Compared with the LCR of *M. musculus* zyxin in the same assay, recruitment is less pronounced, but significant (Fig. 5G), as is true for vertebrate Tes

Taken together, our results are consistent with a model in which actomyosin-mediated tension generated in elongating embryos leads to strain-dependent recruitment of TES-1 to junctions during elongation, stabilizing them against the rigors of mechanical stress during morphogenesis. In this sense, elongating epidermal cells in the *C. elegans* embryo are subject to “self-injury”, as they must remodel their junctional-proximal actin networks during the dramatic change in shape that these cells undergo. It is likely that LIM-domain dependent stabilization of junctional proximal actin filaments is only one component of an apparatus that stabilizes and repairs such filaments. For example, our previous experiments indicated that UNC-94/tropmodulin is recruited to the same junctions, where it presumably protects minus ends of Factin filaments from subunit loss [33]. Recruitment of TES-1 to these same junctions could stabilize CCC-dependent actin networks by allowing strained F-actin at the CCC to self-heal, by recruiting additional F-actin to these networks, or both.

## Abbreviations used in this study

AJ: adherens junction
CCC: cadherin-catenin complex
CFB: circumferential filament bundle
DIC: differential interference contrast
LIM: Lin-l1, Isl-1, Mec-3
PET: Prickle, Espinas, Testin

---

**Supplemental Figure 1.**
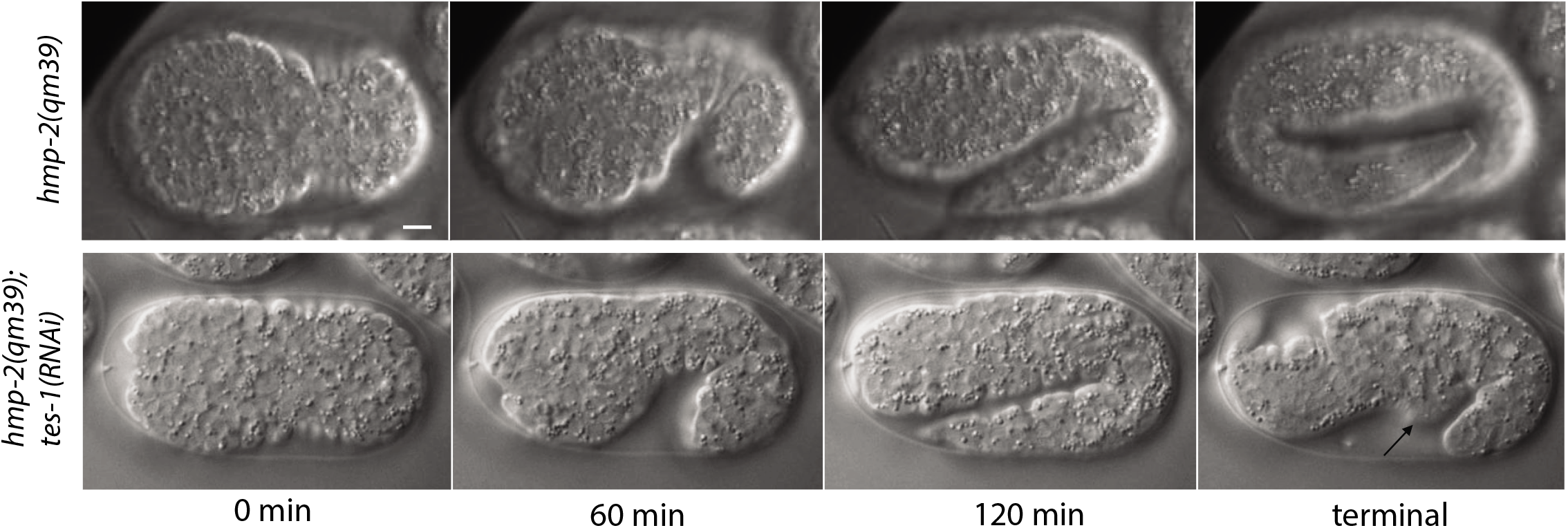
Depletion of TES-1 enhances defects in a *hmp-2* hypomorph. (Top) *hmp-2(qm39)* embryos are viable and display subtle body morphology defects. (Bottom) In *hmp-2(qm39);tes-1(RNAi)* embryos, cells leak out of the ventral midline in terminally arrested embryos (right panel, arrow).

**Supplemental Figure 2.**
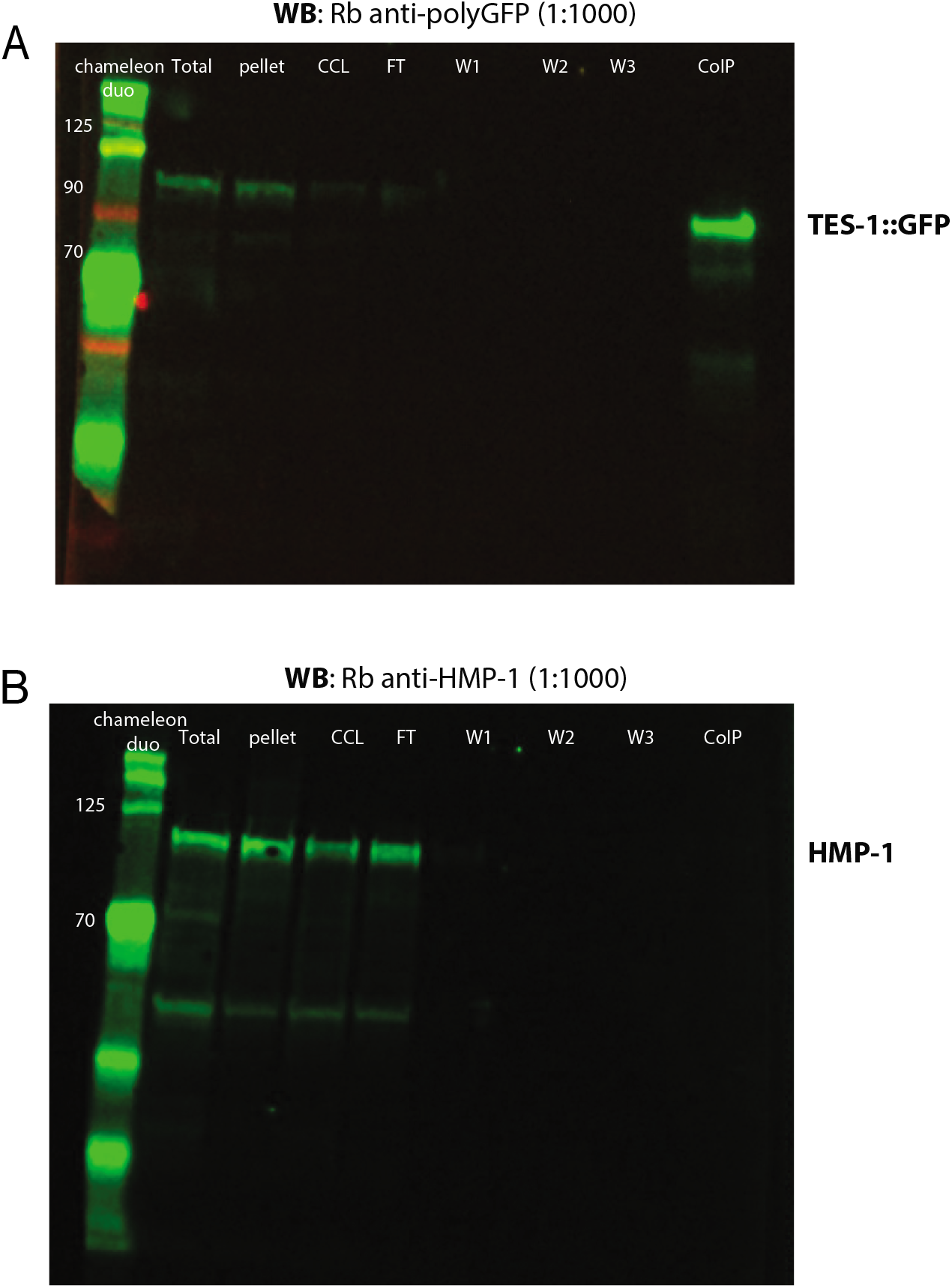
TES-1 cannot coimmunoprecipitate HMP-1/α-catenin. TES-1::GFP was immunoprecipitated from an extract of mixed stage embryos, and the resulting proteins were blotted and probed with anti-GFP and anti-HMP-1 antibodies. TES-1::GFP is substantially enriched in the IP fraction (A), demonstrating that anti-GFP antibodies can coIP TES-1::GFP. Although in a parallel preparation HMP-1 can be detected in the total lysate, pellet and wash fractions, it is undetectable in the IP fraction (B).

**Supplemental Figure 3.**
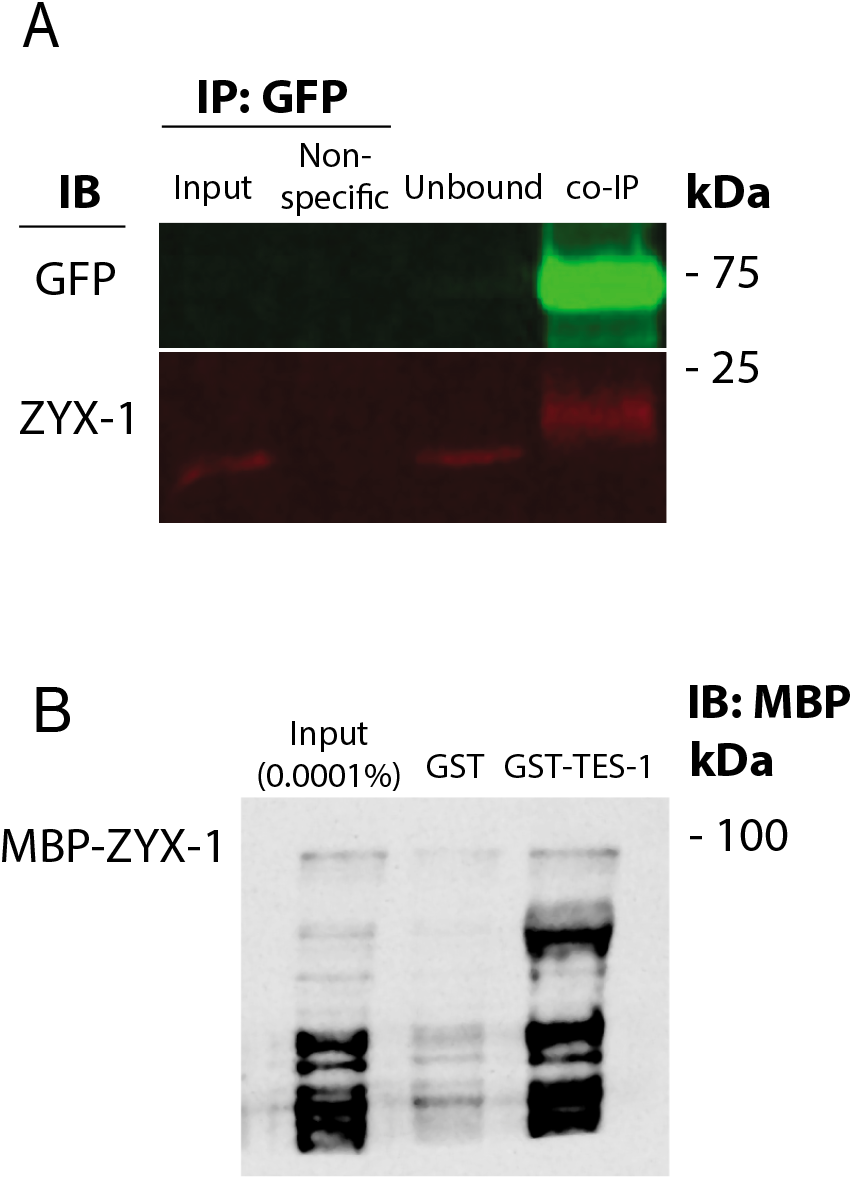
TES-1 only weakly binds ZYX-1/zyxin. (A) Co-immunoprecipitation of TES-1 and ZYX-1. TES-1::GFP was immunoprecipitated from an extract of mixed stage embryos, and the resulting protein was blotted and probed with anti-GFP and anti-ZYX-1 antibodies. ZYX-1 is substantially enriched in the IP fraction. (C) Pulldown using recombinant ZYX-1/zyxin and TES-1/Tes. Extracts of bacteria expressing ZYX-1-MBP were incubated with either GST or GST-TES-1. The resulting mixture was purified using glutathione beads, blotted, and probed using anti-MBP antibodies. MBP-ZYX-1 and TES-1GST weakly interact at substoichiometric levels.

**Supplemental Video 1.** Time lapse movie comparing *hmp-1(fe4)* homozygous and *hmp-1(fe4);tes-1(RNAi)* embryos. Time is shown in hours:minutes. The latter fail consistently during early elongation, and all develop the Humpback phenotype.

## Materials and Methods

### Nematode Strains and Genetics

*C. elegans* strains were maintained using standard methods [48]. Bristol N2 was used as wildtype. The following alleles were utilized in this study. LGI: *hmp-2(qm39); let-502(sb118);* LGII: *zyx-1(gk190); mel-11(it26);* LGIV: *tes-1(ok1036);* LGV: *hmp-1(fe4)*. The following transgenic arrays were made for this study: *tes-1::gfp, tes-1 ΔPET::gfp, tes-1 ΔLIM1::gfp, tes-1 ΔLIM2::gfp, tes-1ΔLIM3::gfp, tes-1 ΔLIM1-3::gfp*. Each of the *tes-1* constructs contained 5kb of endogenous promoter.

### Plasmids

A ~5kb genomic sequence containing 2kb promoter and entire genomic region of *tes-1* was PCR amplified using Phusion polymerase (NEB). The primers used were: 5’ GCGTCGACGAGTTTTTGTCAAGAGTAAGAC and 3’ GCCCCGGGATCAACTGATCATCCGGATTCG. The PCR product was digested with *SalI* and *SmaI* and ligated into a similarly digested Fire lab vector pPD95.75, which contains the GFP sequence. A frameshift was repaired via PCR to generate a *Ptes-1(2kb)::tes-1::gfp* construct (pAML224). To generate *Ptes-1(5kb)::tes-1::gfp*, additional promoter sequence was PCR amplified using Phusion polymerase. The primers used were: 5’ GCCTGCAGGAAGACAACGCTTGTCAAGAAT and 3’ GCGTCGACATTTTGCCCTCGAAATGCAATAC. The PCR product and pAML224 were digested using *PstI* and *SalI* and ligated together to generate pAML224v2. The identity of pAML224v2 was confirmed via sequencing. Domain deletions were performed using circle PCR as described previously [35].

### Microinjection

DNA was microinjected into worms as described previously [49]. Briefly, injection mixes consisting of 5ng/μl of transgenic *tes-1* DNA constructs, 20 ng/μl of junk DNA (F35D3) and 75 ng/μl of *rol-6(su1006)* transgenic marker DNA were microinjected into both gonads of hermaphrodites. Progeny were screened for the presence of *rol-6(su1006)*, and stable lines were established by passaging of worms.

Injection RNA interference was performed as described previously [50]. dsRNA was generated using an Ambion T7 and/or T3 Megascript kits; templates included yk662b10 (*hmr-1*), yk285a2 *(ajm-1)*, yk1054c06 *(zyx-1)*, yk282d2 *(zoo-1)* (NEXTDB, http://nematode.lab.nig.ac.jp/).

### Antibody and Phalloidin Staining

Immunostaining was performed using freeze-cracking [51]. Staining was performed as described previously [52]. Embryos were mounted onto poly-L-lysine-coated ring slides and incubated with primary antibodies in PBST and 5% non-fat dry milk overnight at 4°C. Embryos were then incubated with secondary antibodies in PBST and 5% non-fat dry milk for approximately three hours at room temperature. The following primary antibodies were used: 1:1000 mouse-anti-GFP (Invitrogen), 1:1000 rabbit-anti-GFP, 1:4000 polyclonal rabbit-anti-HMP-1, 1:4000 polyclonal rabbit-anti-HMR-1 and 1:200 monoclonal mouse-anti-AJM-1 (MH27). The following secondary antibodies were used: 1:50 anti-rabbit IgG Texas Red, 1:50 anti-rabbit FITC, 1:50 anti-mouse Texas Red and 1:50 anti-mouse FITC.

Phalloidin staining of mutant and wild-type embryos was used to visualize actin in fixed embryos [27]. Embryos were mounted on poly-L-lysine-coated ring slides and fixed using the following: 4% paraformaldehyde, 0.1 mg/mL lysolecithin, 48 mM Pipes pH 6.8, 25 mM Hepes pH 6.8, 2 mM MgCl2, and 10 mM EGTA for 20 minutes at room temperature. 1:20 Phalloidin-488 was incubated with embryos at room temperature for 90 minutes. Images of stained embryos were acquired as described below.

For co-immunostaining and phalloidin staining, embryos were gathered in a 1.5 mL Eppendorf tube and permeabilized with a solution of 4% paraformaldehyde, 10% Triton-X-100, 48 mM Pipes pH 6.8, 25 mM Hepes pH 6.8, 2 mM MgCl2 and 10mM EGTA for 20 minutes at room temperature. Embryos were incubated overnight in PBST+5% dry milk+1:1000 rabbit-anti-GFP at 4C on a nutator. Secondary antibodies (1:10 Phalloidin-666 and 1:50 anti-rabbit FITC) were incubated for 2 hours at room temperature. Images of stained embryos were acquired as described below.

### Confocal Microscopy

Spinning-disc confocal images were acquired with a Z-slice spacing of 0.2μm for imaging of actin, 0.3 μm for embryos stained for both GFP and actin, and 0.5μm for all other imaging using either Perkin Elmer Ultraview or Micromanager software [53, 54] and a Nikon Eclipse E600 microscope connected to a Yokogawa CSU10 spinning disk scanhead and a Hamamatsu ORCA-ER charge-coupled device (CCD) camera. Junctional/cytoplasmic signal measurements were performed as described previously [55]. Fisher’s exact test calculations were performed online at https://www.socscistatistics.com/tests/fisher/default2.aspx; other statistical analyses were performed using GraphPad Prism v. 9.0 software (GraphPad Software, San Diego, California USA, www.graphpad.com).

### DIC Imaging

Four dimensional DIC movies were gathered on either a Nikon Optiphot-2 connected to a QImaging camera or an Olympus BX5 connected to a Scion camera. Mounts were made as previously described (Raich et al., 1999). ImageJ plugins (http://worms.zoology.wisc.edu/research/4d/4d.html) were used to compress and view movies.

### Protein Expression and Purification

GST- and SUMO-His-tagged proteins were expressed in BL21-Gold(DE3) *Escherichia coli* cells and purified as described [47, 56]. Cells were induced with 0.1mM IPTG at 18°C for 16 hours.

Wash and elution buffers were as follows: GST wash (1X PBS, 500mM NaCl, 0.1% Tween-20, and 1mM DTT), GST elution (50mM Tris pH 8.0, 0.3% glutathione, 150mM NaCl), His wash (50mM Na-Phosphate pH 8.0, 300mM NaCl, 0.1% Tween-20, 10mM Imidazole), and His elution (250mM Imidazole, 100mM NaCl, 10% glycerol, 50mM Hepes pH 7.6). For actin-pelleting assays, the GST tag was cleaved from GST-TES-1 using ProTEV Plus (Promega), according to manufacturer’s instructions.

### Directed Yeast Two-Hybrid Assays

Yeast two-hybrid assays were performed as in [35]. Either full-length HMP-1 or HMP-2 yeast two-hybrid plasmid [57] was transformed into Y2H Gold yeast singly or with a plasmid encoding full-length ZYX-1 [58]. Positive single transformants were tested for autoactivation, and double transformants were patched onto SD/-Leu/-Trp/X-a-gal/AurA plates. Colonies that grew and turned blue were considered positive for a direct interaction.

### Actin-Pelleting Assays

Actin co-sedimentation assays were performed as described previously [47]. Briefly, 5μM purified, cleaved proteins (quantified via a Bradford Assay) were incubated at room temperature for one hour with 0 or 5μM polymerized chicken F-actin (Cytoskeleton, Inc.). BSA was used a negative control, and SUMO-His-HMP-1 [39] was used as a positive control. Samples were then centrifuged at 100,000 rpm for 20 min at 4°C in a TLA-120.1 rotor using a Beckman Optima tabletop ultracentrifuge. Samples were run on 12% SDS-PAGE gels, stained with Coomassie Brilliant Blue, and bands were quantified using ImageJ.

### Co-immunoprecipitations and Western Blots

*C. elegans* expressing TES-1::GFP were grown in liquid culture as previously described [59]. Co-immunoprecipitations were completed as in [33]. Western blots were performed as described previously [60], using rabbit anti-GFP, rabbit anti-HMP-1 [39] and mouse anti-ZYX-1 [58] primary antibodies and Li-COR IRDye^®^ secondary antibodies to detect proteins.

### Stress fiber strain site assay

A *tes-1 LCR::mCherry* construct was designed and expressed using the procedures described in detail in [2]. Briefly, a synthetic gBlock DNA encoding a mammalian codon-optimized version of the LIM1-3 domain of TES-1 was ordered from IDT (Coralville, Iowa) and cloned into a CMV-driven expression vector that fused the C-terminus of LCR(Tes) to mCherry, and used to transfect zyxin ^-/-^ mouse embryo fibroblast cells (MEFs) rescued with stably integrated GFP-zyxin. Transfected MEFs were imaged on an inverted Nikon Ti-E microscope (Nikon, Melville, NY) with a Yokogawa CSU-X confocal scanhead and Zyla 4.2 sCMOS Camera (Andor, Belfast, UK). A 405 nm laser coupled to a Mosaic digital micromirror device (Andor) was used to locally damage stress fibers. Kymography of TES-1(LIM1-3)::GFP was performed using ImageJ as described in [2].

## Acknowledgements

Some strains were provided by the *C. elegans* Genetics Center, which is funded by the NIH Office of Research Infrastructure Programs (P40 OD010440). cDNA clones for *hmr-1, ajm-1, zyx-1, zoo-1, hmp-1, and tes-1* (yk collection) were provided by Yuji Kohara (National Institute of Genetics). AL, BL, SM, and JH were supported by NIH grant R01GM058038 awarded to JH. BM was supported by a Gilliam Fellowship from the Howard Hughes Medical Institute, and by an Advanced Opportunities Fellowship and a COVID-19 dissertation completion fellowship from the University of Wisconsin-Madison. SB and AA were supported by NIH grant R35GM134865 awarded to AA. JW and MG were supported by NIH grant R01GM104032 and Army Research Office Multidisciplinary University Research Initiative W911NF1410403 awarded to MG.

